# Improved metagenome assemblies and taxonomic binning using long-read circular consensus sequence data

**DOI:** 10.1101/026922

**Authors:** J.A. Frank, Y. Pan, A. Tooming-Klunderud, V.G.H. Eijsink, A.C. McHardy, A.J. Nederbragt, P.B. Pope

## Abstract

DNA assembly is a core methodological step in metagenomic pipelines used to study the structure and function within microbial communities. Here we investigate the utility of Pacific Biosciences long and high accuracy circular consensus sequencing (CCS) reads for metagenomics projects. We compared the application and performance of both PacBio CCS and Illumina HiSeq data with assembly and taxonomic binning algorithms using metagenomic samples representing a complex microbial community. Eight SMRT cells produced approximately 94 Mb of CCS reads from a biogas reactor microbiome sample, which averaged 1319 nt in length and 99.7 % accuracy. CCS data assembly generated a comparative number of large contigs greater than 1 kb, to those assembled from a ∼190x larger HiSeq dataset (∼18 Gb) produced from the same sample (i.e approximately 62 % of total contigs). Hybrid assemblies using PacBio CCS and HiSeq contigs produced improvements in assembly statistics, including an increase in the average contig length and number of large contigs. The incorporation of CCS data produced significant enhancements in taxonomic binning and genome reconstruction of two dominant phylotypes, which assembled and binned poorly using HiSeq data alone. Collectively these results illustrate the value of PacBio CCS reads in certain metagenomics applications.

## INTRODUCTION

Metagenome assembly is a key methodological stage in all environmental sequencing projects, which has significant repercussions on all down-stream analyses such as taxonomic classification, genome reconstruction, and functional gene annotation. It is commonly a very complex process, with many sequencing platform-specific issues such as read length and number. Similarly, there are also many sample-specific issues such as the numbers, frequencies, types and sizes of microbial genomes present in highly diverse communities. The goal of metagenomic assemblies is relatively straightforward: obtain large contig sizes coupled with the fewest possible misassemblies. However, metagenomic assemblies often consist of a fragmented collection of short contigs, which are difficult to taxonomically and functionally assign accurately. There are at least two current approaches to metagenomic assembly: (*i*) assembly of all data^1^, which is typically computationally demanding, or (*ii*) using binning or normalization methods to select subsets of reads that are then assembled separately^2, 3^. Methods that use data from multiple sequencing platforms are still infrequent, despite indications that combined approaches yield improvements in contig length and integrity^4^.

Current sequencing technologies offer a range of read lengths. Methods that produce short reads (<250 nucleotides (nt)) such as Illumina can generate high sequencing depth with minimal costs, however when used for analyzing complex communities data assembly typically requires massive computational resources and the resulting contigs remain relatively short^1^. In theory, longer read sequencing technologies can overcome many of the known assembly problems associated with short reads, however these technologies have traditionally been accompanied with one or more inherent shortcomings, such as lower sequencing depth, higher costs and higher error rates. Several technologies exist that can produce longer reads. For example, Ion Torrent and Roche 454 offer read lengths of up to 400 nt and 1000 nt, respectively, but these technologies are more costly per base pair and are vulnerable to generating homopolymer (single-nucleotide repeats) sequencing errors. Pacific Biosciences (PacBio) has designed a sequencing technology based on single-molecule, real-time (SMRT) detection that can provide much greater read lengths, with ∼50% of reads in a single run exceeding 14 kb and 5% exceeding 30 kb^5^. High error rates, reported as high as 15% in individual reads, have previously prevented the use of raw PacBio reads in metagenomics^6,7^. Interestingly, the error rates may be reduced by using circular consensus sequencing (CCS) that entails the repeated sequencing of a circular template, and subsequent generation of a consensus of individual DNA inserts. Consensus quality increases with each sequencing pass, and this approach can ultimately result in high-quality sequences of about 500 to ∼2,500 nt in length with greater than 99% accuracy (Q20 or better)^8,9^.

Here, we present various applications of PacBio CCS data in a metagenomic analysis of the complex microbial community in a commercial biogas reactor. We compare individual assemblies of short read HiSeq2000 and PacBio CCS data as well as hybrid assemblies of subsets from both platforms. PacBio CCS data provides a dramatic improvement in the assembly of universal marker genes in comparison to HiSeq2000 data, allowing for custom training data for phylogenomic binning algorithms and accurate taxonomic binning of assembled contigs from both data types. Subsequently this enabled enhancements in genome reconstructions of uncultured microorganisms that inhabit complex communities.

## RESULTS

### PacBio CCS reads improve assembly statistics

For the purpose of this study we analyzed and compared two sequence datasets generated from the same biological sample, a methanogenic biogas reactor microbiome containing an estimated 480 individual phylotypes, hereafter referred to as Link_ADI (**Table S1**). These datasets comprised approximately one lane of HiSeq sequence data and data from eight PacBio SMRT cells, respectively. HiSeq sequencing entailed 175 nt library construction and generation of 2 × 100 nt paired end sequence data, totaling approximately 149 million read pairs. For PacBio, a library was constructed with inserts of approximately 1.5 kb, which were sequenced using a RS II instrument and P4-C2 chemistry. A total of 522,695 PacBio reads were generated with a mean accuracy of 86 %, totaling approximately 3.3 Gb. Of these reads, 71,254 were CCS that averaged 99.7% accuracy and 1,319 nt in length (totaling 95.4 Mb)(Fig. 1). Given the two different sequencing platforms, multiple assembly algorithms were used. MIRA 4.0^10^ was used to assemble the PacBio CCS reads, which resulted in approximately 46% of the CCS reads assembling into 2,181 contigs averaging 4,459 nt with the max contig length of 65,165 nt (**Table S2**). SOAPdenovo2^11^ was used to assemble 18.5 Gb of HiSeq data generated for Link_ADI, which produced 3,035,577 contigs (average length 189 nt; 55,633 > 1 kb) with a maximum length of 148,797 nt.

**Figure 1.**
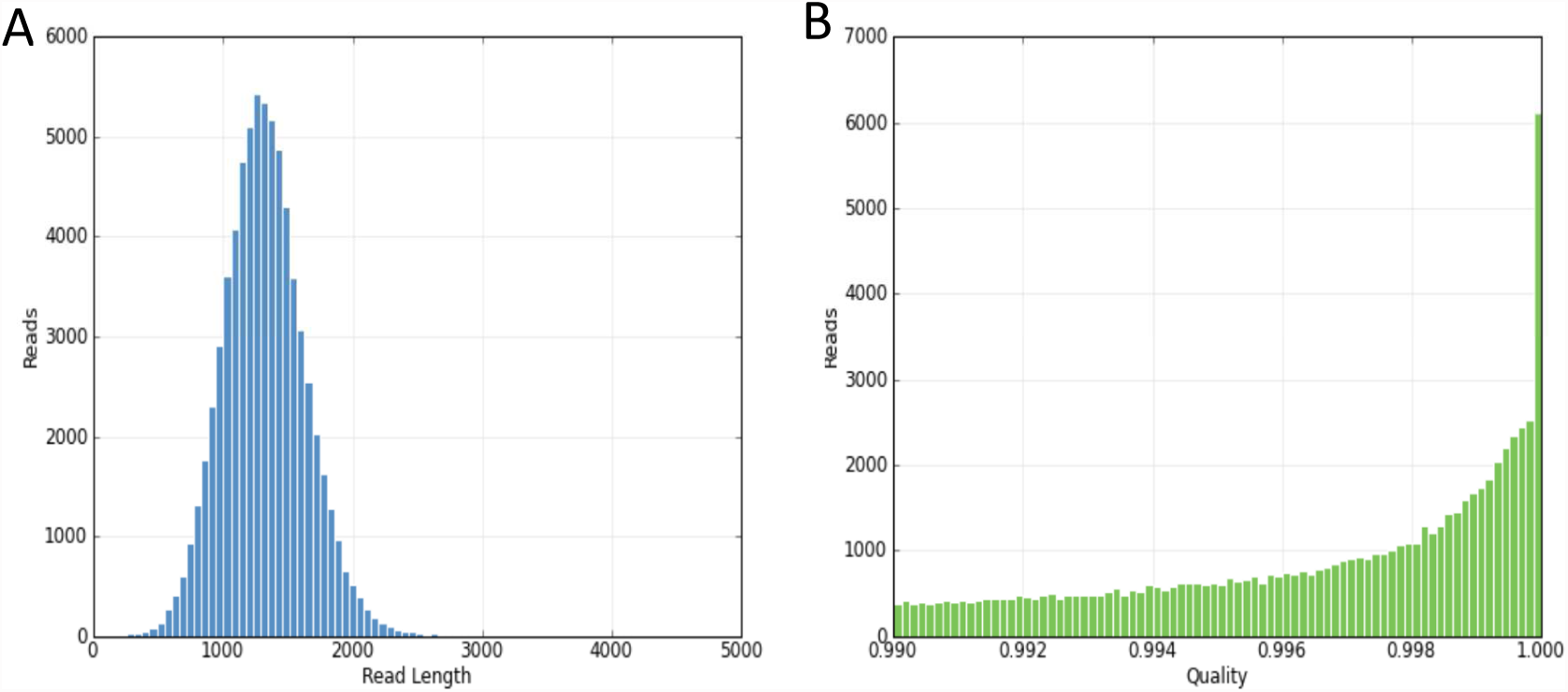
Read length and quality distribution of PacBio “Circular Consensus Sequence” (CCS) reads produced from a Link_ADI-derived shotgun library (∼1.5 kb inserts) sequenced on a PacBio RS II instrument using P4-C2 chemistry. In total, eight SMRT cell were used for sequencing. (**a**) Read length distribution of PacBio CCS reads that passed a 0.99 quality score for which an average of 10 insert passes was required (**b**) Quality distribution of the 71,254 PacBio CCS reads that passed the 0.99 cutoff using the SMRT portal (average 99.7%).

Comparing the statistics from the two assemblies showed that, despite the much smaller size of the raw PacBio CCS dataset (around 190-fold less sequence), the total length of large contigs produced from the MIRA assembly was in the range of those produced from the HiSeq assembly (Fig. 2 and **Table S2**). The MIRA assembly produced 34,513 contigs and unassembled reads that were greater than 1 kb in length, which totaled approximately 54.9 Mb (**Table S2**). In contrast, the HiSeq assembly generated 55,633 contigs greater than 1 kb (134.2 Mb). The total size of the 100 biggest MIRA contigs totaled 52% of the equivalent HiSeq subset. Attempts to perform hybrid assemblies using raw HiSeq and PacBio CCS reads were ultimately unsuccessful, presumably due to the large number of sequencing reads and a paucity of algorithms customized for this particular hybrid input (to our knowledge). Therefore, as an alternative we used a downstream approach that was more amenable to our datasets and available assemblers. Both subsets of assembled HiSeq and CCS contigs greater than 1 kb (including unassembled CCS reads > 1 kb) were further assembled using the “Sanger”-era program CAP3^12^, which was designed for use with long sequencing reads. The resulting hybrid assemblies (Fig. 2 and **Table S2**), which include unassembled contigs from both platforms, provided an increase in mean contig length (PacBio: 1475 nt, HiSeq: 189 nt, Hydrid: 2056 nt) as well as an increase in cumulative nucleotides from contigs larger than 10 kb (PacBio + HiSeq: 21.01 Mb, Hybrid: 26. 8 Mb) and 25 kb (PacBio + HiSeq: 6.5 Mb, Hybrid: 9.3 Mb).

**Figure 2.**
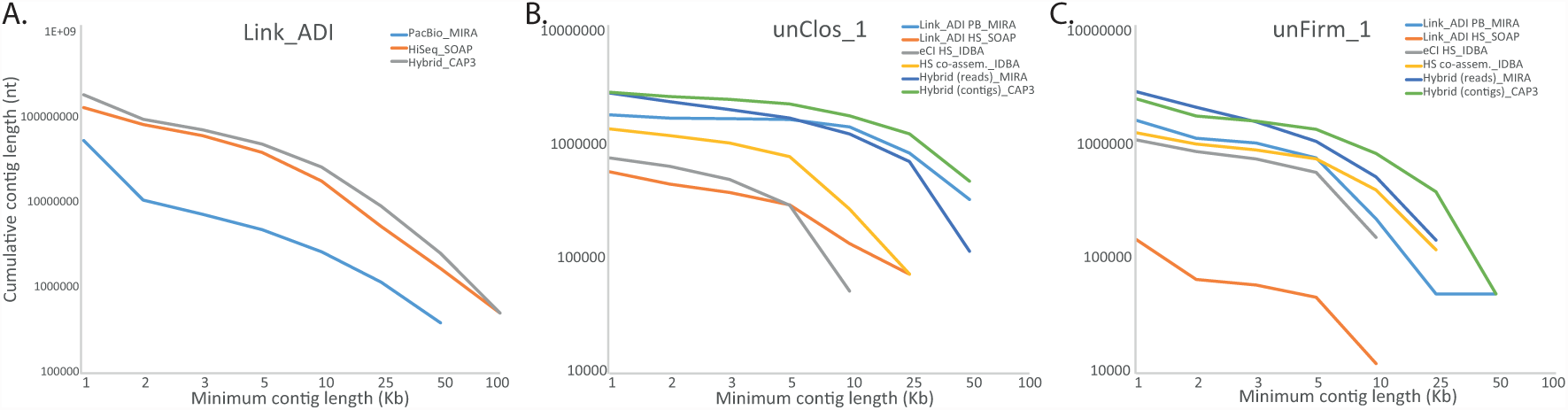
Cumulative number of assembled nucleotides in contigs of different minimum lengths for (**a**) Link_ADI, (**b**) unClos_1, and (**c**) unFirm_1. Each line corresponds to a different sample (Link_ADI or eCI, where noted), sequencing method (HiSeq [HS] or PacBio [PB]), different assembly method (co-assembly across samples Link_ADI and eCI, hybrid using mapped reads from HiSeq and PacBio, or hybrid using contigs from HiSeq and PacBio), or assembly program (CAP3, IDBA_UD, MIRA, or SOAPdenovo).

### PacBio CCS reads improve genome binning of difficult to assemble phylotypes

Community characterization of Link_ADI using short subunit (SSU) rRNA gene amplicon analysis identified approximately 480 individual phylotypes, of which two exhibited high relative abundance and no close taxonomic relationship to cultivated bacterial species (**Table S1**). Phylotype unClos_1 is an as-yet uncultured bacterium affiliated to the Clostridiales family and was estimated to represent ∼36 % of the total microbiome, whereas unFirm_1 is a deeply-branched uncultured representative affiliated to the Firmicutes, accounting for ∼5 %. In order to functionally characterize both phylotypes and determine their contribution to the microbiomes metabolic network, we sought to reconstruct and annotate their genomes. Given the high levels of relative abundance, both organisms were anticipated to be represented by high DNA levels within the metagenomic datasets, and thus conducive to greater assembly in terms of coverage and contig length. First pass comparisons of the assembled HiSeq contigs focusing on contig coverage, size and GC %, gave no clear patterns that are indicative of several numerically dominating organisms (i.e. a cluster of large high-coverage contigs within a narrow GC % range, Fig. 3c). In contrast, coverage vs GC % comparisons of assembled PacBio CCS contigs revealed one clear cluster of higher coverage contigs that were large and within a narrow GC % range (Fig. 3a).

**Figure 3.**
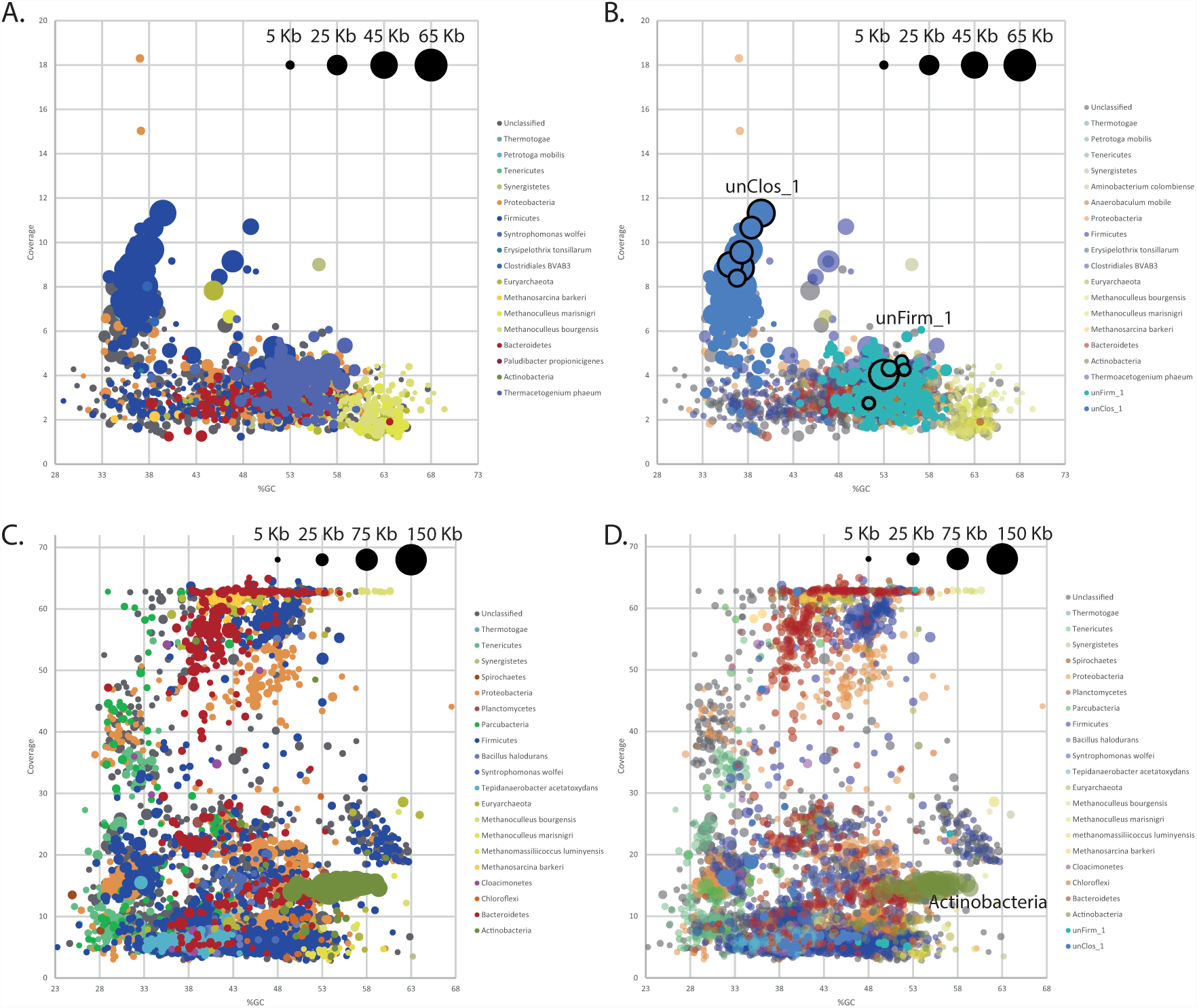
Visualization of GC %, coverage and size of assembled contigs generated from PacBio CCS (**a**, **b**) and HiSeq data (**c**, **d**) from a biogas reactor microbiome (Link_ADI). Contigs are coloured based on taxonomic binning that was performed using PhyloPythiaS+ under default settings (**a**, **c**) and after including custom phylotype-specific training data (**b**, **d**). Contig lengths are indicated by circle sizes. PacBio CCS contigs that contain marker genes and were used as training data for phylotype unClos_1 and unFirm_1 are outlined in black. For the purposes of clarity, only HiSeq contigs greater than 5 kb are represented (**c**, **d**).

Phylogenomic binning methods were subsequently used in attempts to recover genome sequence information for unClos_1 and unFirm_1 and for as many other phylotypes as possible. The presence of only one biological sample and DNA extraction, pre-determined the use of sequence compositional binning algorithms and prevented the use of temporal and/or multi-sample binning methods that have been recently shown to produce accurate genomes from metagenomic datasets^13,14^. PhylopythiaS+^15^ was initially used to assign taxonomy to PacBio CCS and HiSeq contigs (greater than 1 kb), which produced very few taxonomic assignments to a strain or species level (**Table S3**). Instead, the vast majority of contigs were binned to higher-ranking taxa at a phylum or order level, implying that the data provides limited functional and structural insights into the individual organisms making up the microbial community. This result was not unexpected as the SSU rRNA gene analyses indicated that the Link_ADI microbiome is composed of uncharacterized species (**Table S1**) that are distantly related to the available prokaryotic genomes in NCBI used to train PhylopythiaS+.

In cases where PhyloPythiaS and its predecessors have had phylotype-specific training data (at least 100 kb) from a given metagenome, the binning and genome reconstruction of the target phylotype has proven to be highly accurate^16,17^. Therefore, to improve the resolution of PhyloPythiaS+ we compiled as much phylotype-specific training data as possible. All contigs were evaluated for coverage vs. GC% metrics and the presence of taxonomically informative marker genes^18^, with the aim of identifying contigs that correspond to the abundant phylotypes identified in our samples and can therefore be used as training data. The complexity and fragmented nature of the HiSeq assembly (Fig. 3c) made identification of species-specific genome information problematic. This had direct implications on the ability to obtain the ∼100 kb high-confidence assemblages of training data that are required for accurate species level binning^17^. However, the increased length and improved clustering of the assembled PacBio CCS contigs provided large and accurate training data collections for unClos_1 and unFirm_1 in particular. We pooled together six contigs totaling 200 kb for unClos_1 and seven contigs totaling 107 kb for unFirm_1 (Highlighted in Fig. 3b). Interestingly this included large contigs that encoded complete SSU rRNA operons, which are notoriously difficult to assemble using short-read NGS data, such as reads obtained using HiSeq. In total, we identified 17 SSU rRNA gene fragments in the PacBio CCS contigs and 86 when including unassembled reads (compared to six in the HiSeq contigs greater than 1 kb) with three matching unClos_1 (from contigs totaling 96 kb in length).

Both the total collection of HiSeq contigs greater than 1 kb and the PacBio CCS contigs, including unassembled reads, were binned with the custom training model for PhylopythiaS+, that includes all the available prokaryotic genomes in NCBI and the two phylotype-specific contig subsets described above. The output produced a greatly improved recovery of phylotype-level binning for both unClos_1 and unFirm_1 in both HiSeq and PacBio CCS contigs from Link_ADI (Fig. 4). For unClos_1, 189 PacBio sequences (PacBio contigs and unassembled CCS reads, totaling 1,913,759 nt) and 182 HiSeq contigs (600,903 nt) were assigned to the phylotype (**Table S2**). 576 PacBio sequences (1,710,231 nt) and 77 HiSeq contigs (151,790 nt) were binned to unFirm_1. The binning of unClos_1 and unFirm_1 contigs also revealed patterns that indicate assembly differences between PacBio CCS and HiSeq. Despite the indications from the SSU rRNA gene amplicon analyses that phylotypes unClos_1 and unFirm_1 were the most abundant in Link_ADI, neither phylotype were attributed to the longest HiSeq contigs (Fig. 3d). Nine of the ten largest HiSeq contigs from Link_ADI binned to the Order Actinomycetales (Fig. 3c), totaling around 2.2 Mb over 203 contigs (**Table S2**). Only one phylotype affiliated to the Actinomycetales was identified in SSU rRNA gene amplicon analysis, which was ranked 61^th^ most abundant (**Table S1**). In addition, the coverage for each of the Actinomycetales-affiliated HiSeq contigs was on average approximately two-fold higher than the contigs binning as unClos_1 (Fig. 3d). In contrast, the Actinomycetales-affiliated PacBio CCS contigs were much shorter and exhibited lower coverage than unClos_1 (Fig. 3, Fig. 4).

**Figure 4.**
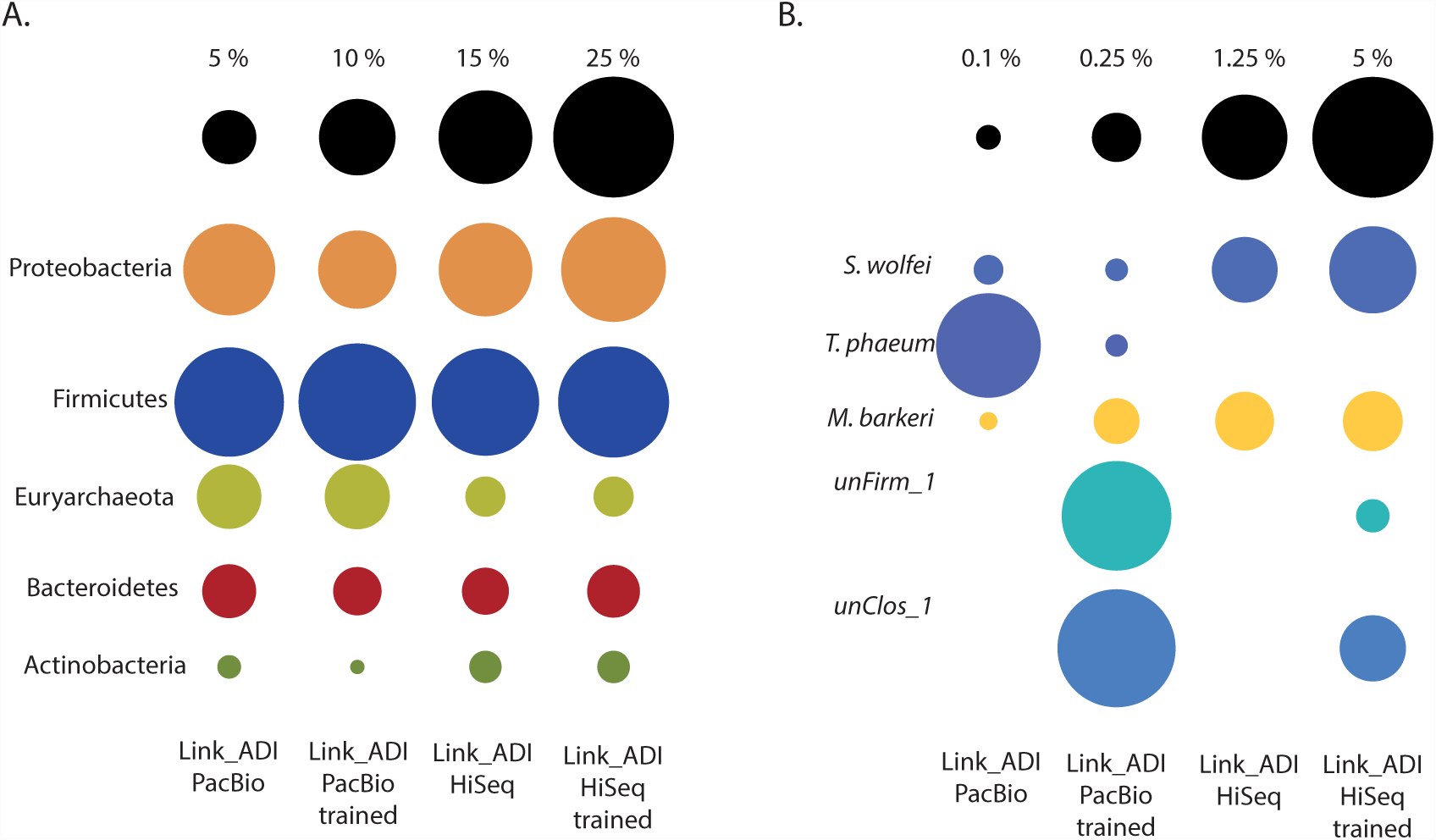
Selected taxonomic bins generated via PhyloPythiaS+ binning using default settings with and without use of custom training data. Circle size indicates relative bin size; for complete binning information see **Table S3**. The proportion of total DNA binned in the major phyla (**a**) represented in the Link_ADI microbiome was similar for both PacBio CCS and HiSeq contigs regardless of the use of training data. However, use of training data enhanced the recovery of unClos_1 and unFirm_1 (**b**) in both the PacBio and HiSeq assemblies. Differences between the sequencing methods were also evident at a species level where some abundant species assembled and binned better with PacBio (*Thermacetogenium phaeum*, unClos_1, and unFirm_1), whereas others produced better results with HiSeq data (*Syntrophomonas wolfei* and *Methanosarcina barkeri*).

The custom trained PhyloPythiaS+ with training data obtained from the PacBio CCS contigs also showed enhanced binning when used for other biological samples and metagenomics datasets where unClos_1 and unFirm_1 were found (Fig. 5). An independently created cellulose enrichment (eCI) was inoculated from Link_ADI and exhibited comparable population structure, with both unClos_1 and unFirm_1 demonstrating numerical dominance (**Table S4**). Similar to the Link_ADI HiSeq dataset, assembly of eCI (IBDA_UD^19^) did not generate long marker-gene encoding contigs representative of unClos_1 and unFirm_1, and phylotype-specific binning was not possible using this dataset alone (Fig. 5a). Therefore, training data generated from the Link_ADI PacBio CCS dataset was used to taxonomically bin the eCI HiSeq dataset (Fig. 5b). The binning produced after training improved cluster visualization, and binning assignments were concurrent with coverage vs GC % comparisons, which indicated explicit clusters for each phylotype (Fig. 5b). Subsequently, the recovery of genomic information linked to the unClos_1 and unFirm_1 phylotypes was substantially larger (**Table S3**). Similar to Link_ADI, assembly discrepancies were also noted in enrichment eCI, where unClos_1 and unFirm_1 were the most abundant organisms (approximately ∼48 % and ∼7 % relative abundance, respectively), but did not assemble into the largest contigs, which again affiliated with the Actinobacteria (Fig. 5).

**Figure 5.**
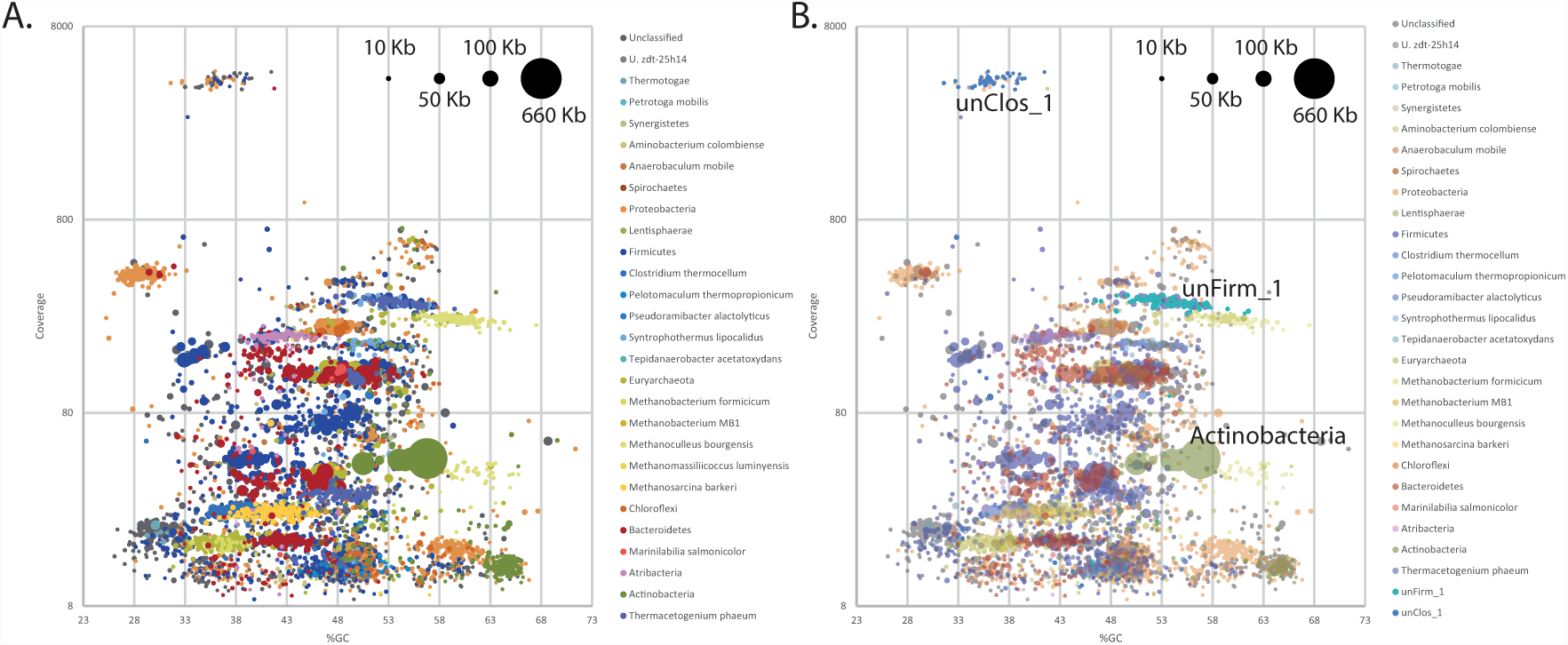
Visualization of GC %, coverage and size of assembled contigs generated from eCI HiSeq data. Sample eCI originated from a lab-scale enrichment grown on cellulose that was inoculated from Link_ADI. Contig lengths are indicated by circle sizes. Contigs are coloured based on phylogenetic binning that was performed using PhyloPythiaS+ under default settings (**a**) and PacBio-derived custom phylotype-specific training data (**b**). For the purposes of clarity, only HiSeq contigs greater than 5 kb are represented.

### Hybrid assembly of genome bins improves overall genomic reconstruction

In an effort to reconstruct improved genomes for both unClos_1 and unFirm_1, we used a two-step hybrid assembly approach that was refined to include only PacBio and HiSeq data that binned to either phylotype. With the intention of generating as complete as possible genomes, we used all genomic material that was available for both phylotypes from both the Link_ADI and eCI samples. Binned HiSeq contigs from Link_ADI and the cellulose enrichment eCI datasets were first deconstructed into individual reads and then pooled into one file prior to assembly using IBDA_UD. These hybrid HiSeq contigs were then assembled together with Pacbio CCS contigs and unassembled reads binned to the same phylotype. This phylotype-specific hybrid approach improved genome reconstruction in terms of total genome size as well as improved average contig length and large contig assembly (Fig. 2b-c and **Table S2)**. For unClos_1, a total of 1178 sequences (PacBio contigs, unincorporated PacBio reads, and co-assembled Link_ADI and eCI HiSeq contigs) 3,350,596 nt in length were assembled into 430 contigs (and unincorporated sequences) greater than 1 Kb totaling 3,030,306 nt. For unFirm_1, 1,212 sequences (3,037,687 nt) from unFirm_1 were assembled into 815 contigs greater than 1 Kb, totaling 2,650,713 nt. Hybrid MIRA assemblies that used the individual sequencing reads (that formed the original contigs) instead of a two-step approach using CAP3, resulted in contigs that were on average smaller for both unClos_1 and unFirm_1 (Fig. 2b-c and **Table S2**).

## DISCUSSION

Many of the commonly used second generation sequencing methods in (meta)genome sequencing provide gigabases of data. While this provides high levels of sequencing depth per sample, the short read lengths can restrict the ability to assemble longer contigs, particularly when evaluating complex microbial communities. Specific exemplary problems include the presence of genes with low evolutionary divergence between organisms or repetitive genomic regions that are larger than a sequencing read (e.g., rRNA operons). One way of circumventing this is by combining multiple sequencing technologies that can overcome each other’s limitations. For example, Illumina HiSeq provides high sequencing depth, but with low sequencing breadth; in other words this technique has a high ability to sample across multiple genomes with the drawback that individual reads sample a very small proportion of each genome. This can be complemented by additional PacBio sequencing, which has high breadth (providing at least 10-30-fold more data per read), but a lot lower depth. By combining the two methods, one has a higher probability of covering regions problematic for short read sequencing methods. Several studies have illustrated this convincingly for bacterial genomes, where a hybrid Illumina-PacBio approach has enabled near-complete chromosome closure with no necessary secondary sequencing or primer-walking methods^20^. Previously, the high error rate of PacBio reads (∼86%) has prevented their use in metagenomic analysis of complex communities, where the coverage required to compensate the erroneous reads was not financially or technically feasible. However, use of the CCS provides high quality long reads that are suitable for metagenomic applications. Here we illustrate the features that PacBio CCS data may bring to a metagenomics project, with respect to increased contig lengths, assembly of problematic genomic regions, improved phylogenomic binning, and genome reconstruction of the uncultured phylotypes that dominate microbial communities.

Specific benefits of the PacBio CCS contigs for Link_ADI were the considerably larger average contig sizes as well as the number of large contigs, with the later being comparable to the HiSeq assembly that was generated from 190-fold more data. In metagenomic analyses, larger contigs are key to producing higher quality output that is needed for downstream applications such as taxonomic assignments^17^, gene calling, and annotation of operons that often exceed 10 kb in length^16^. The assembly output from both platforms varied considerably in both contig size and distribution (Fig.2,Fig. 4 and **Table S2**). In particular, numerically dominating organisms did not necessary assemble into the largest HiSeq contigs, irrespective of species diversity or the assembly algorithms used (Fig. 3b,Fig. 3d and Fig. 5), which in contrast transpired for PacBio CCS contigs (Fig. 3a-b). Despite the similar size of the PacBio CCS and HiSeq > 1 kb contig datasets available for binning, the size of the unClos_1 and unFirm_1 genomic bins obtained from the PacBio CCS data were, on average, ∼3x and ∼6x larger, respectively (Fig. 4 and **Table S2**). Another observation was the examples of PacBio CCS contigs containing difficult to assemble regions such as SSU rDNA. On average, PacBio CCS contigs that contained relevant SSU rDNA data were 15-fold larger than the SSU rDNA containing HiSeq contigs. Conventional composition-based binning was shown to be substantially improved with the addition of PacBio-derived custom training data that contained genomic information specific for unClos_1 and unFirm_1 (Fig. 4 and **Table S3**). The collection of these phylotype-specific training subsets was only possible in the PacBio CCS contig dataset, since neither phylotype produced contigs of sufficient length in HiSeq datasets. Hence, this approach presents an alternative means to reconstruct genomes in instances were phylotypes are not conducive to HiSeq assembly and experimental design that will not allow multiple sample timepoints or several differential DNA extractions, which are necessary for accurate binning algorithms that use differential coverage of populations^13,14^.

Whilst this study shows the potential value PacBio CCS reads can exert upon a metagenomics study, there is certainly room for improvement. One of the key concerns with the use of PacBio CCS reads is data wastage with respect to the number of reads generated and the number that pass CCS quality cutoffs. One may expect that upcoming PacBio upgrades and increased read lengths will produce a higher amount of high-quality CCS reads and thus less wastage. Notably, closer examination reveals that read wastage is also applicable for the use of Illumina in metagenomic applications. For example, in the present study only 35.6% of the paired-end HiSeq reads assembled into contigs greater than 1,000 nt, an arbitrary cutoff that is used in many metagenomic analyses.

Hybrid assemblies for both the total community dataset and phylotype-specific bins produced improvements (Fig. 2 and **Table S2**), and this represents just a start. In the future, there will be access to better long read data and it is anticipated that further improvement of assembly algorithms customized to incorporate multiple sequencing technology inputs will improve hybrid assembly performance. Regardless, these aspects need further attention in moving forward, so that the full potential of longer read technology can be exploited to deepen our insight into complex microbial communities. This study also shows that as long reads become more common, they will make further software extensions of binning algorithms such as PhyloPythiaS+ very valuable and will allow automatic assignment of training contigs to novel phylotypes and not just the higher ranking assignments. Increased capabilities to reconstruct accurate genomes representative of uncultured microorganisms are of major importance since they allow accurate mapping of community metabolism and are a prerequisite for meaningful “meta-omic” studies that may reveal genes and/or proteins with novel functions that cannot be recognized by bioinformatics alone.

## METHODS

### Samples

Sample Link_ADI was obtained from a commercial biogas reactor in Linköping, Sweden, fed on a mixture of slaughterhouse waste, food waste, and plant biomass (Reactor I)^21^. Sample eCI was taken from a batch enrichment using the same commercial biogas plant as inoculum source and cellulose as substrate^22^.

### DNA extraction and sequencing

Total genomic DNA was prepared using the FastDNA Spin Kit for Soil (MP Biomedicals, Santa Ana, CA, USA). For both Link_ADI and cEI, an aliquot of 200 μl was used for DNA extraction following the manufacturer’s protocol. For SSU rRNA gene sequencing, library preparation was performed as per manufacturers recommendations (Illumina, 2013). V3 and V4 regions of bacterial SSU rRNA genes were amplified using the 341F (5’-<u>TCGTCGGCAGCGTCAGATGTGTATAAGAGACAG</u>CCTACGGGNGGCWG CAG-3’) and 785R (5’-<u>GTCTCGTGGGCTCGGAGATGTGTATAAGAGACAG</u>GACTACH VGGGTATCTAATCC-3’) modified primer set^23^, where the underlined sequence corresponds to the Illumina adaptor. The amplicon PCR reaction mixture (25 μl) consisted of 12.5 ng microbial gDNA, 12.5 μl iProof HF DNA polymerase mix (BioRad) and 0.2 μM of each primer. The PCR reaction was performed with an initial denaturation step at 98°C for 30 s, followed by 25 cycles of denaturation at 98°C for 30 s, annealing at 55°C for 30 s, and extension at 72°C for 30 s followed by a final elongation at 72°C for 5 min. A new PCR reaction was carried out to attach unique 6 nt indices (Nextera XT Index Kit) to the Illumina sequencing adaptors to allow multiplexing of samples. The PCR conditions were as follows: 98°C for 3 min., 8 cycles of 95°C for 30s., 55°C for 30 s., and 72°C for 30 °C, followed by a final elongation step at 72°C for 5 min. AMPure XP beads were used to purify the resulting 16S rRNA amplicons. The 16S rRNA amplicons were quantified (Quant-IT™ dsDNA HSAssay Kit and Qubit™ fluorometer, Invitrogen, Carlsbad, CA, USA), normalized and then pooled in equimolar concentrations. The mulitiplexed library pool was then spiked with 25 % PhiX control to improve base calling during sequencing. A final concentration of 8 pM denatured DNA was sequenced on an Illumina MiSeq instrument using the MiSeq reagent v3 kit chemistry with paired end, 2 × 300 bp cycle run. HiSeq Shotgun sequencing runs were performed on libraries (175 nt, to ensure overlap and allow for merging of the paired-ends) prepared from Link_ADI and enrichment cEI DNA using TruSeq PE Cluster Kit v3-cBot-HS sequencing kit (Illumina Inc.). In addition, libraries prepared from Link_ADI DNA were shotgun sequenced using the PacBio RS II Single Molecule, Real-Time (SMRT®) DNA Sequencing System. Library. The library was prepared using the PacBio 2 kb library preparation protocol and sequenced on 8 SMRT cells using P4-C2 chemistry.

### SSU rRNA gene amplicon analysis

Paired end reads were joined using the QIIME v1.8.0 toolkit included python script join_paired_ends.py (with the default method fastq-join) and quality filtered (at Phred >=Q20) before proceeding with downstream analysis^24^. USEARCH61 was used for detection of chimeric sequences followed by clustering (at 97% sequence similarity) of non-chimera sequences and denovo picking of OTUs^25,26^. Joined reads were assigned to OTUs using the QIIME v1.8.0 toolkit^24^, where uclust^27^ was applied to search sequences against a subset of the Greengenes database^28^ filtered at 97% identity. Sequences were assigned to OTUs based on their best hit to the Greengenes database, with a cut-off at 97% sequence identity. Taxonomy was assigned to each sequence by accepting the Greengenes taxonomy string of the best matching Greengenes sequence. filter_otus_from_otu_table.py (included with QIIME) was used to filter out OTUs making up less than 0.005% of the total using default parameters and --min_count_fraction set to 0.00005 as previously reported^29^.

### Raw data assembly

HiSeq data from Link_ADI was assembled using SOAPdenovo-63mer (SOAPdenovo2 http://soap.genomics.org.cn/soapdenovo.html) using the following the parameters: -K 51 -p 40 setting max_rd_len=125, avg_ins=100, reverse_seq=0, and asm_flags=1. PacBio reads for Link_ADI were filtered using the SMRT portal, with only those CCS reads that produced a minimum accuracy of 0.99 (average 10 passes) being considered for further analysis (ranging from one to three kb in length). PacBio CCS reads were assembled using slightly modified parameters in MIRA 4.0 (http://sourceforge.net/p/mira-assembler/wiki/Home/): COMMON_SETTINGS -DI:trt=./-NW:cmrl=warn \PCBIOHQ_SETTINGS -CL:pec=yes. Sequence data from enrichment cEI was trimmed using sickle pe (version 0.940 https://github.com/najoshi/sickle) with default parameters, converted to an interleaved FASTA using the program fq2fa (bundled with IDBA_UD) with the parameters --merge --filter, and assembled with IDBA_UD v1.1.1 (http://i.cs.hku.hk/∼alse/hkubrg/projects/idba_ud/index.html) using the parameters -- pre_correction --num_threads 15 --maxk 60.

### Identification of marker genes in contigs

For the identification of protein coding marker genes, open reading frame calling was first performed using MetaGeneMark^30^ version 1 metagenome ORF calling model (gmhmmp -m MetaGeneMark_v1.mod -f G -a -d). Output was subsequently converted into a multiple FASTA using the included aa_from_gff.pl script. The resulting proteins sequences were compared against the 31 AMPHORA marker gene HMMs using HMMSCAN (part of HMMER version 3.0^31^), that form the basis of an automated phylogenomic inference pipeline for bacterial sequences^18^. The marker genes used are: *dnaG*, *frr*, *infC*, *nusA*, *pgk*, *pyrG*, *rplA*, *rplB*, *rplC*, *rplD*, *rplE*, *rplF*, *rplK*, *rplL*, *rplM*, *rplN*, *rplP*, *rplS*, *rplT*, *rpmA*, *rpoB*, *rpsB*, *rpsC*, *rpsE*, *rpsI*, *rpsJ*, *rpsK*, *rpsM*, *rpsS*, *smpB* and *tsf*. Matches with e-values of < 1.e^-5^ were considered legitimate. SSU rDNA searches were conducted using BLASTN (-e 1e-20 -r 1 -q -1 -v 5 -b 5 -F F) against a database of phylogenetically diverse representative sequences from sequenced genomes^32^.

### Genomic binning

The GC % was calculated for each contig and the coverage values for each were provided by each assembler (IDBA_UD provides a single coverage value, MIRA provides average coverage, and SOAPdenovo provides k-mer coverage). From this, we created a table of GC % versus coverage for each contig, allowing us to visualize clustering of contigs. Using contig clustering and marker gene analysis of our PacBio contigs (because they are on average longer and contain greater marker gene representation including SSU rDNA fragments), we were able to generate phylotype-specific training data for the two most abundant organisms (unClos_1 and unFirm_1). These subsets consisted of contigs totaling more than 100 kb, the minimum necessary for custom binning using PhyloPythiaS+^15^. Contigs that met the criteria for phylotype-specific training data were larger than 7 kb, exhibited consist coverage (+- 2x) and GC% (+- 3%) values and encoded a SSU rRNA gene or marker gene that demonstrated phylogenomic grouping with the representative OTU sequence identified via 16S rRNA gene amplicon analysis. Binning was performed using PhyloPythiaS+ using both default settings, against a database consisting of all publically available prokaryotic genomes in NCBI, and with our custom training data.

### Co- and hybrid assembly

Various merged assemblies were performed in an attempt to improve assembly statistics of the Link_ADI community metagenome and the genome reconstructions of dominate phylotypes (unClos_1 and unFirm_1). Hybrid assemblies of whole community contigs (>1 kb) from both the HiSeq and PacBio CCS contig subsets were performed using CAP3^12^ (version date 12/21/07) with default parameters except a minimum overlap percent identity (-p) of 0.95.

In order to reconstruct as large as possible genomes for unClos_1 and unFirm_1, we performed hybrid assemblies of binned contigs for each phylotype from all of our samples including the PacBio and HiSeq data from Link_ADI and the HiSeq data from enrichment eCI. This was carried out in two stages. The first stage consisted of mapping HiSeq reads to their corresponding phylotype contigs using BWA mem^33^ (version 0.7.8-r455) with default parameters. The reads that mapped from each sample (Link_ADI and eCI) were identified by parsing the resulting SAM files, pooled together for each phylotype, and co-assembled with IDBA_UD using the same workflow as eCI above into cross-sample HiSeq contigs. The second stage consisted of pooling together the cross-sample HiSeq contigs with the phylotype-specific PacBio contigs, which were hybrid assembled using CAP3, with the same parameters as above. The unincorporated contigs from the hybrid assemblies (contigs that went into the assembly but were not incorporated into hybrid contigs) were also included in the final reconstructed genomes used in this study.

A hybrid assembly of raw sequences between both platforms was also performed using MIRA 4.0. The cross-sample HiSeq reads used above in each co-assembly were used as input along with PacBio reads that mapped to each species-specific bin (identified through the MIRA supplied CAF result file). MIRA 4.0 was run using the following parameters: COMMON_SETTINGS -SK:mmhr=1 -NW:cac=warn -NW:cdrn=no -NW:cmrl=warn \ PCBIOHQ_SETTINGS -CL:pec=yes \ SOLEXA_SETTINGS -CL:pec=yes. For the HiSeq readgroup, the following information was supplied: template_size = 100 400 and segmet_naming = solexa.

## ACKNOWLEDGEMENTS

JAF and PBP are supported by a grant from the European Research Council (336355- MicroDE). The sequencing service was provided by the Norwegian Sequencing Centre (www.sequencing.uio.no), a national technology platform hosted by the University of Oslo and supported by the “Functional Genomics” and “Infrastructure” programs of the Research Council of Norway and the Southeastern Regional Health Authorities. DNA preparations from Biogas reactor samples were supplied by Professor Anna Schnürer and Li Sun from the Department of Microbiology, Swedish University of Agricultural Science, Uppsala, Sweden. SSU rDNA analyses were preformed by Live H. Hagen from Department of Chemistry, Biotechnology and Food Science, Norwegian University of Life Sciences, Ås, Norway. We thank Professor Abigail A. Salyers from the University of Illinois for her helpful advice and correspondence.

## AUTHOR CONTRIBUTIONS

PBP, AJN and VGHE proposed this project. JAF, AJN, ACM and PBP designed the experiments and supervised the project. JAF, ATK and YP did the experiments. JAF, ATK, YP and PBP analyzed the data. JAF, AJN, VGHE and PBP contributed to analysis of the results and paper writing.

## ADDITIONAL INFORMATION

Datasets are available at the NCBI Sequence Read Archive under the BioProject PRJNA294734. The authors declare there is no competing interest. Correspondence and requests for materials should be addressed to Phillip B. Pope (") and Jeremy A. Frank.

